# Female lower urinary tract microbiota do not correspond to IC/PBS symptoms: a case-controlled study

**DOI:** 10.1101/517557

**Authors:** L. Bresler, T.K. Price, M. Tulke, E.E. Hilt, C. Joyce, C.M. Fitzgerald, A.J. Wolfe

## Abstract

**ABSTRACT:** *OBJECTIVE:* Current etiology of interstitial cystitis/painful bladder syndrome (IC/PBS) is poorly understood and multifactorial. Recent studies suggest the female urinary microbiota (FUM) contribute to IC/PBS symptoms. This study was designed to determine if the FUM, analyzed using mid-stream voided urine samples, differs between IC/PBS patients and controls.

*MATERIALS AND METHODS:* This prospective case-controlled study compared the voided FUM of women with symptoms of urinary frequency, urgency, and bladder pain for greater than 6 months to the voided FUM of healthy female controls without pain. Bacterial identification was performed using 16S rRNA gene sequencing and EQUC, a validated enhanced urine culture approach. Urotype was defined by a genus present at >50% relative abundance. If no genus was present above this threshold, the urotype was classified as ‘mixed’. Chi-square and Fisher’s exact tests were performed. *P*-values <0.05 were considered significant.

*RESULTS:* A mid-stream voided specimen was collected from 21 IC/PBS patients and 20 asymptomatic controls. Both groups had similar demographics. Urotypes did not differ between cohorts as assessed by either EQUC or 16S rRNA gene sequencing. We detected no significant differences between cohorts in terms of alpha-diversity. Cohorts also were not distinct using Principle Component Analysis or hierarchical clustering. Detection by EQUC of bacterial species considered uropathogenic was high in both cohorts, but detection of these uropathogenic species did not differ between groups (*p*=0.10).

*CONCLUSIONS:* Enhanced culture and modern DNA sequencing methods provide evidence that IC/PBS symptoms may not be related to differences in the FUM, at least not its bacterial components. Future larger studies are needed to confirm this preliminary finding.

## INTRODUCTION

Interstitial Cystitis/Painful Bladder Syndrome (IC/PBS) affects nearly 7.9 million US adult women^1^. The AUA and SUFU define IC/PBS as “an unpleasant sensation (pain, pressure, discomfort) perceived to be related to the urinary bladder, associated with lower urinary tract symptoms of more than six weeks duration, in the absence of infection or other identifiable causes”^2^.

The etiology of IC/PBS is multifactorial, poorly understood, and assumes that the female urinary tract is sterile in the absence of clinical infection. However, our research team and others have shown that the female urinary tract is not sterile; it possesses communities of microbes called the female urinary microbiota (FUM)^3 4 5 6^. Furthermore, the FUM is associated with various lower urinary tract symptoms^7 8 9 10 11 12^. These findings present a new avenue for studying the etiologies of IC/PBS.

Few groups have reported on the FUM of IC/PBS patients. The few existing studies used various urine collection methods and bacterial detection methods, in general had small sample sizes and, therefore, the data conflict^13 14 15^. Our prospective case-controlled study was designed to determine if the FUM of women with and without IC/PBS differs using mid-stream voided urine specimens to avoid pain provocation and analyzed them with expanded techniques.

## MATERIALS AND METHODS

### Study Design and Patient Population

Following Institutional Review Board (IRB) approval, we enrolled 41 female patients; 21 with IC/PBS (i.e., IC Cohort) and 20 without (i.e., Control Cohort). Women in the IC Cohort experienced symptoms of urinary frequency, urgency, and bladder pain for greater than 6 months, meeting the AUA IC/PBS definition. For both cohorts, basic demographics were collected. The members of the IC cohort completed validated questionnaires including the-Beck Anxiety Inventory, Beck Depression Inventory, the Pain Disability Index, the Female Genitourinary Pain Index, the Pain Catastrophizing Scale, the IC Symptom Index Score and the IC Problem Index Score.

### Sample Collection

Midstream voided urine specimens were collected from both cohorts. A vaginal swab specimen was collected from the IC Cohort. A portion of each urine sample was placed in a BD Vacutainer^®^ Plus C&S Preservative Tube for culturing. A separate portion for 16S rRNA gene sequencing was placed at 4°C for less than 4 hours following collection; 10% AssayAssure (Sierra Molecular, Incline Village, NV) was added before storage at −80°C. Puritan Opti-Tranz^®^ Liquid Stuart Swabs were used to collect two aerobic vaginal swab specimens. Each swab was vortexed and diluted in 1ml phosphate buffered saline (PBS). One aliquot was used for culture and one was stored for 16S rRNA gene sequencing, as described above.

### Urine Culture Protocols

A variation of the Expanded Quantitative Urine Culture (EQUC) protocol^6^ was used to culture the biological specimens. 10μL of urine sample or vaginal swab aliquot was spread quantitatively onto BAP, Chocolate, and Colistin Naladixic Acid (CNA) agars (BD BBL™ Prepared Plated Media) and incubated in 5% CO_2_ at 35°C for 48 hours; onto BAP incubated aerobically at 35°C for 48 hours; and onto CDC Anaerobic 5% sheep blood (Anaerobic BAP) agar (BD BBL™ Prepared Plated Media) incubated anaerobically at 35°C for 48 hours. The vaginal swab aliquots were also plated on Thayer-Martin media and incubated in 5% CO_2_ at 35°C for 48 hours. Each distinct colony morphology was sub-cultured at 48 hours to obtain pure culture for microbial identification. Microbial identification was determined using a Matrix-Assisted Laser Desorption/Ionization-Time-of-Flight Mass Spectrometer (MALDI-TOF MS, Bruker Daltonics, Billerica, MA).

### DNA isolation and 16S Sequencing

DNA isolation, polymerase chain reaction (PCR) amplification, and 16S rRNA gene sequencing of urine cultures have been described previously^7^. To avoid contamination, isolation of DNA was performed in a laminar flow hood. Genomic DNA was extracted from 1 ml of urine or 500ul of the vaginal swab aliquot, using previously validated protocols developed for the Human Microbiome Project^16 6 7^. To isolate genomic DNA from these samples, this protocol includes the addition of mutanolysin and lysozyme to ensure robust lysis of Gram-positive and Gram-negative species^16^. Briefly, the aliquoted specimen was centrifuged at 13,500 rpm for 10 min, and the resulting pellet was resuspended in 200 μl of filter-sterilized buffer consisting of 20 mM Tris-Cl (pH 8), 2 mM EDTA, 1.2% Triton X-100, and 20 μg/ml lysozyme and supplemented with 30 μl of filter-sterilized mutanolysin (5,000 U/ml; Sigma-Aldrich, St. Louis, MO). The mixture was incubated for 1 h at 37°C, and the lysates were processed through the DNeasy blood and tissue kit (Qiagen, Valencia, CA), according to the manufacturer’s protocol. The DNA was eluted into 50 μl of buffer AE, pH 8.0, and stored at −20° C.

The hyper-variable region 4 (V4) of the bacterial 16S rRNA gene was amplified via a two-step PCR protocol, as described previously^6 7^. Briefly, in the first amplification, the V4 region was amplified using Illumina MiSeq modified universal primers 515F and 806R. Extraction negative controls (no urine or swab suspension) and PCR-negative controls (no template) were included to assess the contribution of extraneous DNA from reagents. Ten-microliter aliquots of each reaction mixture were run on a 1% agarose gel. Samples containing a band of approximately 360 bp were considered PCR-positive and subjected to further library preparation. Samples with no visible amplified product were considered PCR-negative and not processed further. The PCR-positive reaction mixtures were diluted 1:50 and amplified for an additional 10 cycles, using primers encoding the required adapter sequences for Illumina MiSeq sequencing and an 8-nucleotide sample index. The PCR reaction was purified and size selected using Agencourt AMPure XP-PCR magnetic beads (Beckman Coulter, Pasadena, CA). Each sample was quantified using the Qubit fluorometeric system (Thermo-Fisher, Waltham, MA). The samples were pooled, quantified to a standard volume, and placed in the 2 × 250 bp sequencing reagent cartridge, according to the manufacturer’s instructions (Illumina, San Diego, CA).

Sample barcodes and sequencing primers were removed using the Illumina proprietary MiSeq post-sequencing software. The mothur program (v1.37.4) was used to process the raw sequences by following the recommended MiSeq standard operating procedure^17^. Briefly, mothur produced 16S contigs by combining the paired end reads based on overlapping nucleotides in the sequence reads; contigs of incorrect length for the V4 region (<290 bp, >300 bp) and/or contigs containing ambiguous bases were removed. Chimeric sequences were removed using UCHIME within the mothur package^18^. Subsampling at a depth of 5000 sequences was performed to correct for different sequencing depth of each sample. The sequences were clustered into species-level operational taxonomic units (OTUs) with identity cutoff at 97%^19^. The OTUs were classified using RDP classifier (v2.11) at the genus level^19^. Specimens designated as “undetectable” had <1000 total sequence reads.

### Statistical Analyses

Continuous variables were reported as means with standard deviations (SD); categorical variables were reported as frequencies and percentages. Pearson Chi-square tests (or Fisher’s Exact Tests, when necessary) and 2-sample t-tests (or Wilcoxon Rank Sum tests, when appropriate) were used to compare demographics and culture results (e.g., abundance and diversity) between cohorts. All statistical analyses were conducted using SAS software v9.4 (SAS Institute, Cary, NC) or SYSTAT software version 13.1 (SYSTAT Software Inc., Chicago, IL).

## RESULTS

### Demographics

**Table 1** displays the demographic characteristics of the two cohorts (IC and Control). The entire population had a mean age of 49 and was predominately White/Caucasian (70%) consistent with our clinical patient’s population. The two groups had similar demographics and percentage of EQUC and 16S sequence-positive specimens (*p*>0.05 for all comparisons except vaginal parity).

**Table 1.** Demographic and Clinical Variables for the IC and Control Cohorts.

| Patient and Clinical Variables | Total Cohort (N = 41) | IC Cohort (N = 21) | Control Cohort (N=20) | p-value |
| --- | --- | --- | --- | --- |
| Age (years), mean (SD) | 49 (13) | 50 (13) | 48 (12) | 0.36<sup>a</sup> |
| Race: (N=40) |  | (N=20) |  |  |
| -White/Caucasian | 29 (71%) | 18 (90%) | 11 (55%) | 0.06<sup>b</sup> |
| -Black/African American | 6 (15%) | 1 (5%) | 5 (25%) | 0.09<sup>b</sup> |
| -NHPI | 0 (0%) | 0 (0%) | 0 (0%) | - |
| -Asian | 0 (0%) | 0 (0%) | 0 (0%) | - |
| -Other | 5 (12%) | 1 (5%) | 4 (20%) | 0.18<sup>b</sup> |
| Ethnicity: (N=40) |  | (N=20) |  |  |
| -Hispanic/Latina | 5 (13%) | 3 (15%) | 2 (10%) | 1.00<sup>b</sup> |
| -Not Hispanic/Latina | 35 (87%) | 17 (85%) | 18 (90%) |  |
| BMI (kg/m<sup>2</sup>), mean (SD) | 26.80 (6.40) | 25.52 (5.89) | 28.14 (6.79) | 0.42<sup>a</sup> |
| Vaginal Parity, mean (SD) (N=38) | 2 (1) | (N=20)<br>1 (1) | (N=18)<br>2 (2) | 0.01<sup>a</sup> |
| Menopausal Status: |  |  |  |  |
| -Pre-menopausal | 18 (44%) | 10 (48%) | 8 (40%) | 0.62 |
| -Post-menopausal | 20 (49%) | 10 (48%) | 10 (50%) | 0.88 |
| -Not Sure | 3 (7%) | 1 (4%) | 2 (10%) | 0.61<sup>b</sup> |
| Marital Status |  |  |  |  |
| -Single | 7 (17%) | 5 (25%) | 2 (10%) | 0.41<sup>b</sup> |
| -Married | 29 (71%) | 14 (67%) | 15 (75%) | 0.56 |
| -Divorced | 2 (5%) | 1 (4%) | 1 (5%) | 1.00<sup>b</sup> |
| -Widowed | 3 (7%) | 1 (4%) | 2 (10%) | 0.61<sup>b</sup> |
| Use of Hormone Replacement Therapy | 7 (17%) | 5 (24%) | 2 (10%) | 0.41<sup>b</sup> |
| Smoking | 3 (7%) | 0 (0%) | 3 (15%) | 0.11<sup>b</sup> |
| Alcohol Consumption (N=40) | 21 (53%) | 9 (43%) | (N=19)<br>12 (63%) | 0.20 |
| Use of Antimicrobial Soap (N=39) | 13 (33%) | (N=20)<br>5 (25%) | (N=19)<br>8 (42%) | 0.26 |
| Urine Dipstick: |  |  |  |  |
| - White Blood Cells (WBC) | 0 (0%) | 0 (0%) | 0 (0%) | - |
| - Nitrites | 0 (0%) | 0 (0%) | 0 (0%) | - |
| - Red Blood Cells (RBC) | 0 (0%) | 0 (0%) | 0 (0%) | - |
| EQU C-Positive: |  |  |  |  |
| -Voided Urine (N=38) | 37 (97%) | (N=18) 17 (94%) | 20 (100%) | 0.47<sup>b</sup> |
| -Vaginal Swab (IC Cohort only) (N=18) | 18 (100%) | (N=18) 18 (100%) | - | - |
| 16S Sequencing-Positive: |  |  |  |  |
| -Voided Urine (N=40) | 39 (98%) | 20 (95%) | (N=19) 19 (100%) | 1.00<sup>b</sup> |
| -Vaginal Swab (IC Cohort only) (N=20) | 20 (100%) | (N=20) 20 (100%) | - | - |

Chi-Square Test used unless otherwise indicated. SD = standard deviation.
a – Independent T test, b – Fisher's exact test

### Description of the Lower Urinary Tract and Vaginal Microbiomes in IC Patients

16S rRNA gene sequencing was performed on all (21) IC voided urine and vaginal swab specimens. The majority of voided urine specimens of IC patients had a *Lactobacillus* (11/21; 53%) or Mixed (6/21; 29%) urotype, a measure of bacterial community structure as determined by 16S rRNA gene sequencing (**Supplemental Figure 1**). For most IC patients, the urotype matched the dominant taxa present in the paired vaginal swab specimen (19/21; 90%) (**Supplemental Figure** 1). The specimen types (IC vaginal swab, IC urine, control urine) did not significantly differ by several mean alpha diversity measures that report on the richness, evenness and abundance of community members (**Supplemental Figure 2**).

**Table 2** displays the demographics and validated questionnaire results of the IC cohort subdivided by *Lactobacillus* versus non-*Lactobacillus* urotype. IC Patients with a *Lactobacillus* urotype were younger (*p*=0.01) and more likely to be pre-menopausal (*p*=0.03) than IC patients with a non-*Lactobacillus* urotype, which were more likely to be post-menopausal (*p*=0.009); neither the *Lactobacillus* urotype nor the non-*Lactobacillus* urotypes had significantly different mean scores for any of the validated clinical questionnaires.

**Table 2.**
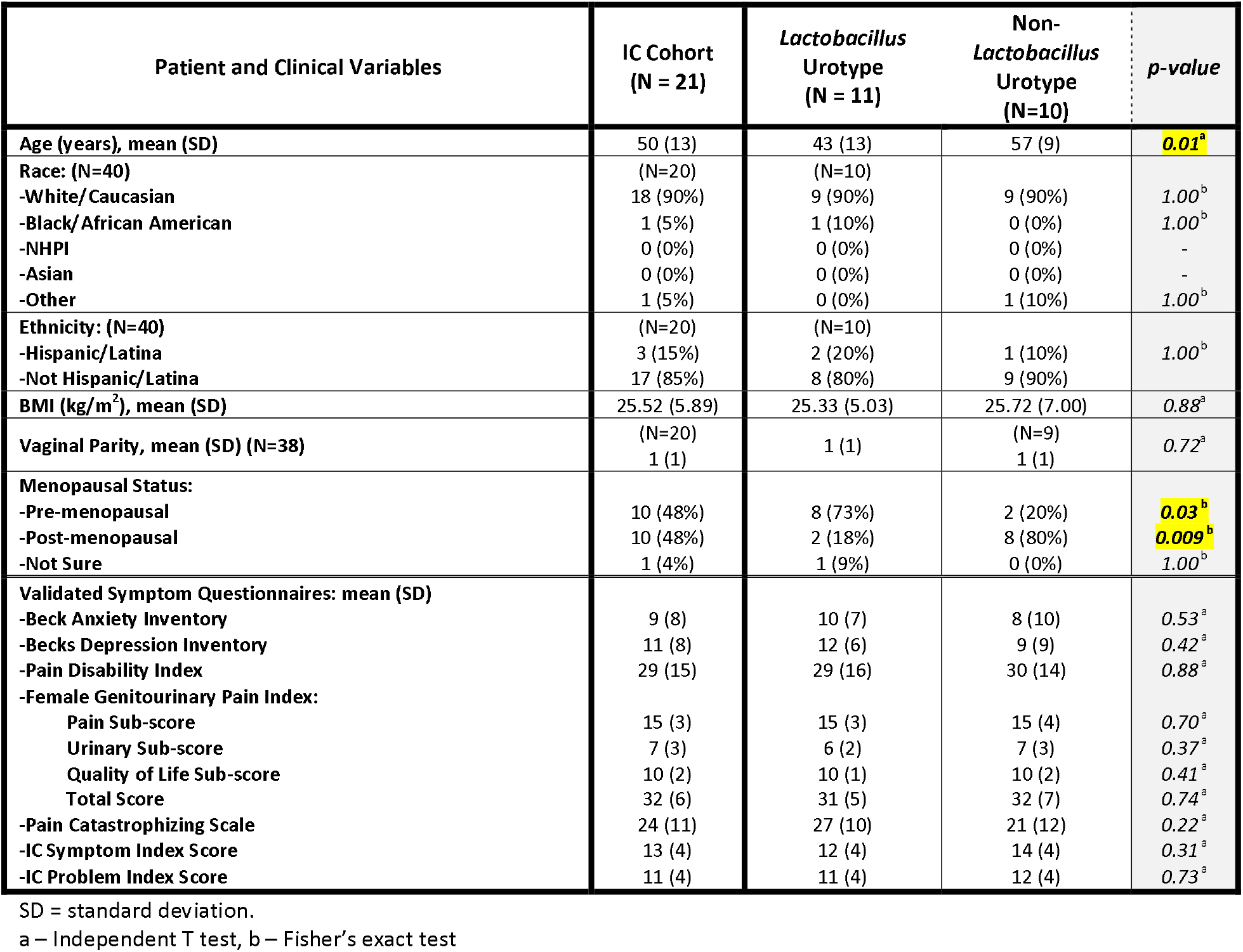
Demographic, Clinical Variables, and Symptom Questionnaire Results for IC Patients with *Lactobacillus* and non-*Lactobacillus* Urotypes determined by 16S rRNA Gene Sequencing.

### Comparison of Lower Urinary Tract Microbiome of IC and Control Patients

16S rRNA gene sequencing was performed on voided urines from 19 of 20 Control specimens. Consistent with the IC Cohort, the Control patients had predominately *Lactobacillus* (9/19; 47%) or Mixed (6/19; 32%) urotypes by 16S rRNA gene sequencing. Mean alpha diversity measures did not differ between the voided urine specimens of the IC and Control cohorts (**Supplemental Figure 2**). Principle component analysis did not show separation between the cohorts (**Figure 1**). A hierarchical cluster analysis of specimens classified at the OTU-level from both cohorts is shown in **Figure 2**, which does not reveal any clear apparent clustering of specimen types or cohort-specific specimens.

**FIGURE 1.**
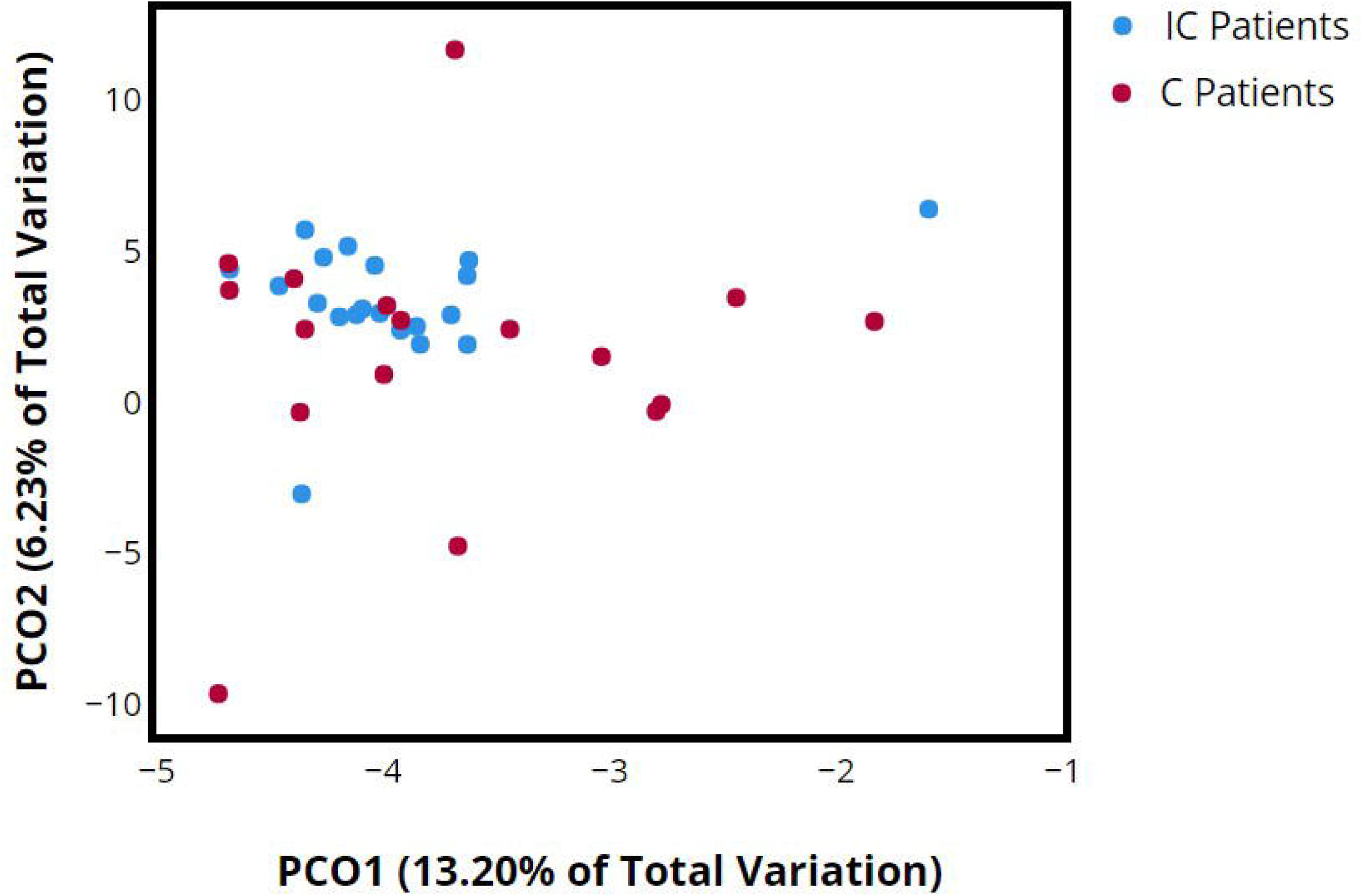
Principle Component Analysis of Mid-Stream Voided Urine Specimens from IC and Control Patients. Principle component analysis comparing 16S rRNA gene sequence data between IC patients (21), blue, and Control patients (C patients) (19), red.

**FIGURE 2.**
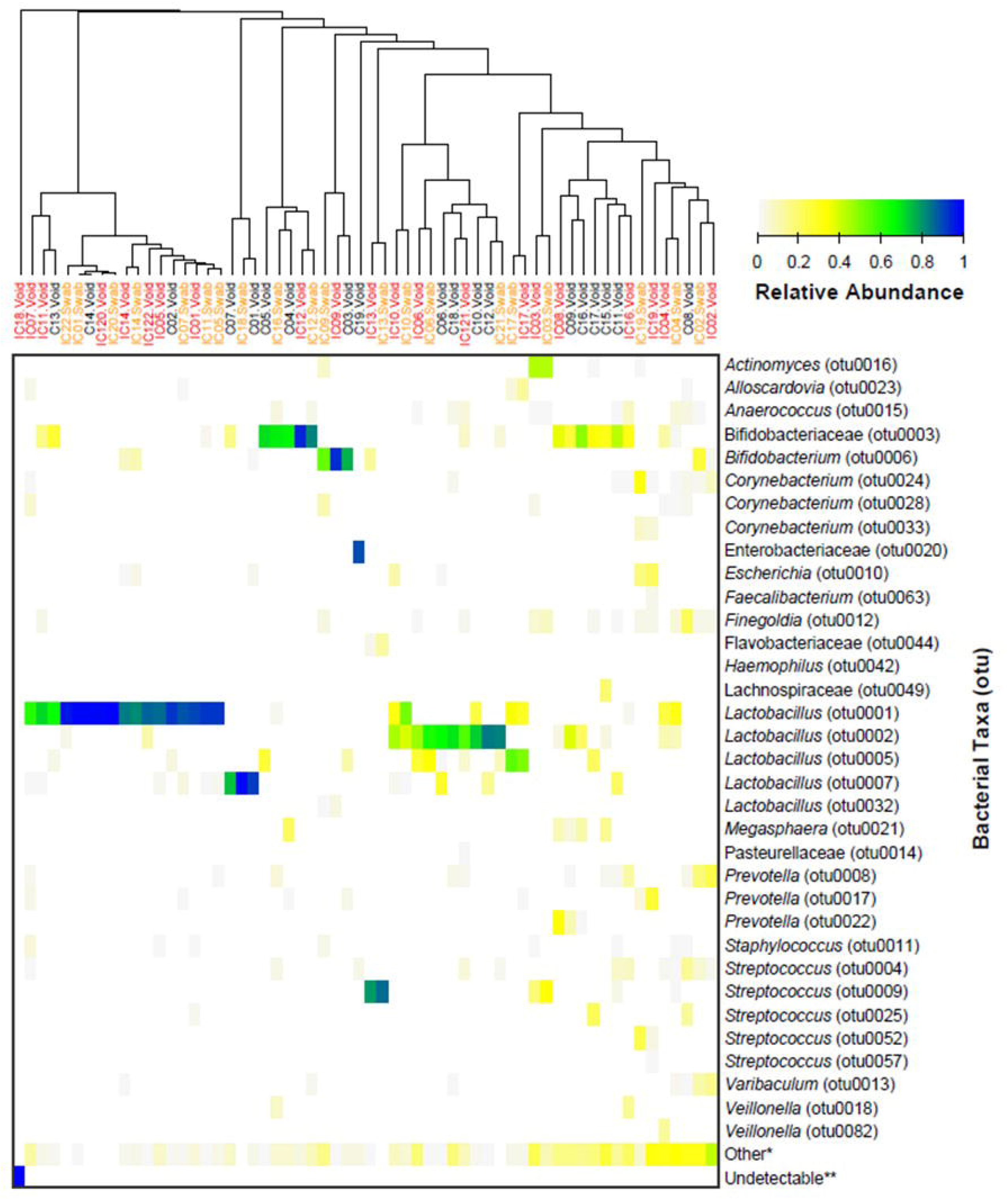
Heatmap of Relative Abundance Values for Common Bacterial OTUs among Cohorts and Specimen Types. Heatmap of the relative abundance of 16S rRNA gene sequence data classified by otu’s. The y-axis lists common bacterial otu classifications in alphabetical order. The x-axis describes the specimen type and patient cohort of the corresponding sample. Specimens listed in red are mid-stream voided urine specimens from IC patients, orange are vaginal swab specimens from IC patients, and black are mid-stream voided urine specimens from Control (C) patients. Data are grouped by hierarchical clustering using the corresponding dendogram.

### Detection of Uropathogenic Bacteria between Cohorts

EQUC was performed on all Control (20) and 18 of the 21 IC urine specimens. Using EQUC, we detected 39 and 51 unique species in the IC (N=18) and Control (N=20) cohorts, respectively. Frequency of detection of *Staphylococcus lugdunensis* (*p*=0.04) and *Streptococcus agalactiae* (i.e. Group B *Streptococcus*) (*p*=0.04) were statistically significant (**Table 3**). Detection of *Escherichia coli* was higher in the IC cohort but did not reach significance (*p*=0.08). Detection of bacterial species typically considered uropathogenic was high in both groups (IC=15/18, 83%; C=20/20, 100%), but not significantly different (*p*=0.10) (**Table 3**).

**Table 3.** List of Uropathogenic Bacterial Taxa Identified in Mid-Stream Voided Urine Specimens of IC and Control patients using EQUC.

| Uropathogenic Bacterial Taxa | IC Cohort<br>(N = 18) | Control Cohort<br>(N=20) | p-value |
| --- | --- | --- | --- |
| *Actinotignum schaalii* | 0 (0%) | 1 (5%) | 1.00<sup>a</sup> |
| *Aerococcus sp.* | 1 (6%) | 5 (25%) | 0.18<sup>a</sup> |
| *Alloscardovia omnicolens* | 3 (17%) | 2 (10%) | 0.65<sup>a</sup> |
| *Candida sp.* | 1 (6%) | 2 (10%) | 1.00<sup>a</sup> |
| *Corynebacterium riegelii* | 0 (0%) | 2 (10%) | 0.49<sup>a</sup> |
| *Enterococcus faecalis* | 9 (50%) | 7 (35%) | 0.35 |
| *Escherichia coli* | 5 (28%) | 1 (5%) | 0.08<sup>a</sup> |
| *Haemophilus influenzae* | 1 (6%) | 0 (0%) | 0.47<sup>a</sup> |
| *Klebsiella pneumoniae* | 0 (0%) | 1 (5%) | 1.00<sup>a</sup> |
| *Neisseria sp.* | 0 (0%) | 1 (5%) | 1.00<sup>a</sup> |
| *Oligella urethralis* | 0 (0%) | 1 (5%) | 1.00<sup>a</sup> |
| *Staphylococcus aureus* | 2 (11%) | 0 (0%) | 0.22<sup>a</sup> |
| *Staphylococcus lugdunensis* | 0 (0%) | 5 (25%) | 0.04<sup>a</sup> |
| *Streptococcus agalactiae* | 1 (6%) | 7 (35%) | 0.04<sup>a</sup> |
| *Streptococcus anginosus* | 9 (50%) | 11 (55%) | 0.76 |

Chi-Square Test used unless otherwise indicated.
a – Fisher's exact test

## DISCUSSION

This study detected no significant differences in the voided FUM of women with and without IC/PBS. Despite the study limitation of small sample size, our numbers were large enough to suggest that microbiota of the lower urinary tract may not contribute to the symptoms in women meeting the clinical definition of IC/PBS. This contradicts recent work by others, who have argued for a link between clinical symptoms of IC/PBS and the FUM^13 14 15^. Abernathy *et al*. hypothesized a protective role of a more diverse and *Lactobacillus*-dominant microbiome. Catheterized urine of women with IC were found to have fewer OTUs and less likely to contain *Lactobacillus* species, particularly *L. acidophilus*. Furthermore, Abernathy *et al*. found that the presence of *Lactobacillus* was associated with improved scores on two IC-specific symptom severity indices, suggesting that the urinary microbiome may influence lower urinary tract symptoms. Although our collection techniques differed and our expanded culture technique additive, these findings were not reproduced in our study nor were the differences in *Lactobacillus* dominance indicative of symptom burden. Siddiqui *et al*. found that *Lactobacillus* predominance was associated with IC/PBS patients (i.e., the opposite trend) using clean catch voided urine specimens. Finally, Nickel *et al*. analyzed a larger cohort of IC/PBS participants (n=233) using clean catch urine during flare vs non-flare pain states and showed no difference in species composition. However, this study did indicate that the IC group had a higher prevalence of fungi. Fungi were not directly analyzed in our cohort. However, we did detect *Candida* species in both cohorts using EQUC.

Strengths of this study were the use of two complementary identification methods: sequencing and culture, as well as the use of vaginal swabs as a comparative specimen to voided urine in the IC cohort. Voided specimens were chosen intentionally so as not to create an IC pain flare by obtaining a catheterized specimen. Furthermore, contemporary urological guidelines do not require a catheterized urine specimen as a part of the diagnostic IC work up (AUA guideline) and not routinely performed in this patient population due to catheterization discomfort and poor patient’s compliance^2^. Limitations include small sample size for both cohorts, lack of vaginal swabs in the controls, and lack of other clinical data in the control group. Although, we did not have an *a priori* sample size estimation, our cohort sizes were similar to those of the Abernethy *et al*. study, which did show differences in the urinary microbiome between groups, so we felt our sample size was adequate for this preliminary analysis. We agree with Nickel *et al*. that the voided urine does not represent the bladder microbiome in our study, but rather the genitourinary tract. Additionally, we also recognize that we did not control antibiotic exposure, similar to Abernethy *et al*.

This case-controlled study of the FUM in predominantly middle-aged women with IC/PBS compared to controls without pain showed no significant differences in the voided FUM between groups. These findings suggest that microbes may not directly contribute to IC/PBS unlike previously reported literature. Larger scale studies using complementary microbial detection techniques similar to ours and assessing multiple urine collection methods (voided and catheterized) would contribute to deeper understanding of the FUM as a potential etiology in IC/PBS.

## Notes

**Funding:** R01: NIH grant from NIDDK awarded to Alan J. Wolfe (R01DK104718) Grant from ICA (Interstitial Cystitis Association) awarded to Colleen Fitzgerald and Larissa Bresler

