## Supplementary figures and images for "Female lower urinary tract microbiota do not correspond to IC/PBS symptoms: a case-controlled study"

### Supplemental Figure 1

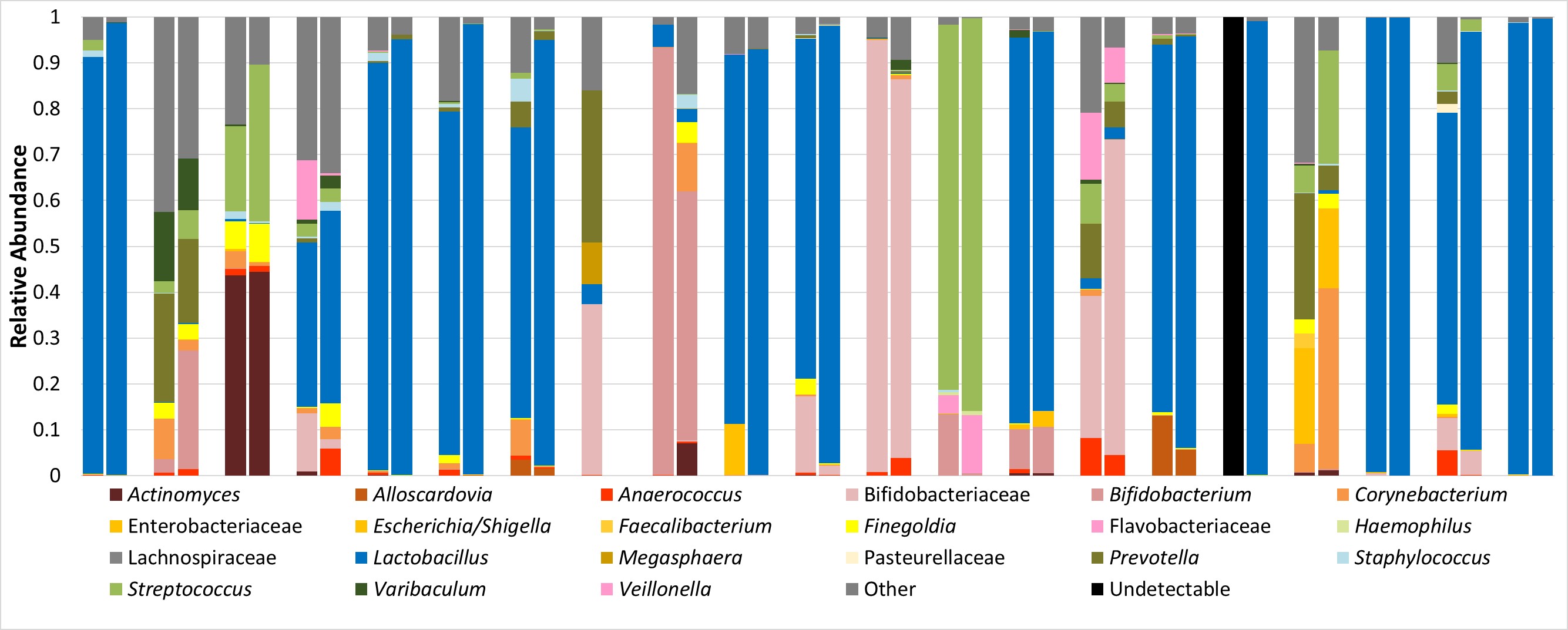

### Supplemental Figure 2

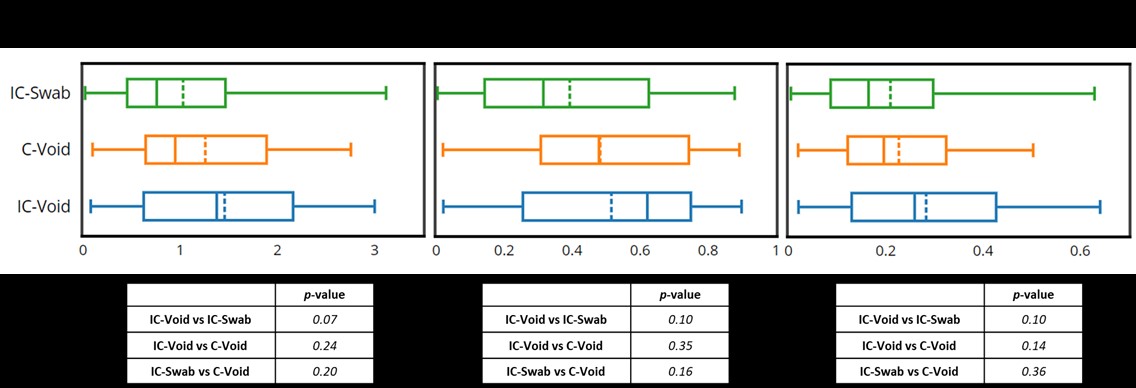
